# Environmental stress determines the colonization and impact of an endophytic fungus on invasive knotweed

**DOI:** 10.1101/2021.09.10.459653

**Authors:** Sigisfredo Garnica, Zhiyong Liao, Samuel Hamard, Frank Waller, Madalin Parepa, Oliver Bossdorf

**Author notes:** Correspondence: Oliver Bossdorf.

## Abstract

1. There is increasing evidence that microbes play a key role in some plant invasions. A diverse and widespread but little understood group of plant-associated microbes are the fungal root endophytes of the order Sebacinales. They are associated with exotic populations of invasive knotweed (*Reynoutria* ssp.) in Europe, but their effects on the invaders are unknown.
2. We used the recently isolated Sebacinales root endophyte S*erendipita herbamans* to experimentally inoculate invasive knotweed and study root colonisation and effects on knotweed growth under different environmental conditions. We verified the inoculation success and fungal colonisation through immunofluorescence microscopy and qPCR.
3. We found that *S. herbamans* strongly colonized invasive knotweed in low-nutrient and shade environments, but much less under drought or benign conditions. At low nutrients, the endophyte had a positive effect on plant growth, whereas the opposite was true under shaded conditions.
4. *Synthesis.* Our study demonstrates that the root endophyte *S. herbamans* has the potential to colonize invasive knotweed fine roots and impact its growth, and it could thus also play a role in natural populations. Our results also show that effects of fungal endophytes on plants can be strongly environment-dependent, and may only be visible under stressful environmental conditions.

## Introduction

Fungal endophytes are a phylogenetically diverse and widespread group of plant-associated microbes (Rodriguez et al., 2009). They can influence the growth and reproduction of individual plants, or their resistance to abiotic stress or natural enemies (Cosme et al., 2016; Kivlin et al., 2013; Mayerhofer et al., 2013; Oberhofer et al., 2014; Rho et al., 2018; Rodriguez et al., 2008). Some of the positive effects are related to the ability of endophytes to improve the nutrition of their host plants (Behie & Bidochka, 2014). There is also evidence that endophytes can influence the diversity and composition of entire plant communities (Afkhami & Strauss, 2016; Aguilar-Trigueros & Rillig, 2016; Clay & Holah, 1999; Rudgers et al., 2004; Rudgers et al., 2005) as well as their associated ecological networks (e.g. herbivores and their parasitoids; Omacini et al., 2001). However, so far our understanding of fungal endophytes is based on experiments with very few taxa, in particular the genus *Neotyphodium* and its asexual stage *Epichloë*: Other fungal systems have been hardly studied, mainly because most fungal endophytes are often difficult to cultivate and thus controlled experiments for testing their ecological functions have so far been impossible.

An important group of fungal endophytes for which this has long been true is the Serendipitaceae family in the order of Sebacinales that contains many species with broad geographic and host ranges (Garnica et al., 2016; Weiss et al., 2011). Previous experimental work has so far been largely restricted to *Serendipita indica* (*Piriformospora indica*), and it showed that *P. indica* stimulates plant growth and influences plant nutrition and tolerances to biotic and abiotic stresses (Achatz et al., 2010; Barazani et al., 2005; Gill et al., 2016; Waller et al., 2005). Our group in Tu◻bingen recently isolated and cultivated another widespread Serendipitaceae species, *Serendipita herbamans*, which is abundant and associated with a broad range of host species and habitats in Central Europe (Riess et al., 2014).

Soil microbes can influence plant growth and stress tolerance, and these effects are to some extent host plant-specific. As a consequence, plant-microbe interactions play a role in structuring plant communities, and there is increasing evidence that they are also important in the invasion of exotic plant species (Callaway et al., 2004; Dawson & Schrama, 2016; Inderjit & van der Putten, 2010; Klironomos, 2002). In general, plant-associated microbes may have positive or negative feedbacks on plants (Bever et al., 2012; van der Putten et al., 2013). If exotic plants accumulate biota with overall more positive effects, maybe because some of the their native pathogens did not make it to the introduced range, this may give invaders an advantage over native plants (Callaway et al., 2011; Maron et al., 2014; Mitchell & Power, 2003; Reinhart et al., 2003). Alternatively, exotic plants may influence soil biota to the detriment of the native plants, e.g. through increasing abundances of their pathogens (Mangla & Callaway, 2007) or disrupting interactions with mutualists (Meinhardt & Gehring, 2012; Stinson et al., 2006).

Most previous research on plant-microbe interactions and plant invasion has focused on soil-borne microbes rather than endophytes, even though fungal endophytes are clearly abundant and diverse also in invasive plant populations (Clay et al., 2016; Shipunov et al., 2008). Besides an interesting series of studies by Aschehoug et al. (2012, 2014) who demonstrated that the leaf endophyte *Alternaria alternata* makes invasive knapweed (*Centaurea stoebe*) more competitive and allelopathic against native North American grasses, there has so far been little experimental work on fungal endophytes and invasive plants.

One of the most problematic plant invaders of temperate Europe and North America is the Japanese knotweed (*Reynoutria japonica*) and its hybrid *R. × bohemica*. Their aggressive growth can damage buildings and other structures, and it has huge impacts on native plant communities and ecosystems (Aguilera et al., 2010; Gerber et al., 2008; Hejda et al., 2009). Because of these ecological and economic costs, there is great interest in controlling invasive knotweed, and in understanding the biological mechanisms contributing to its success. Previous experimental research indicates that chemical or microbial processes belowground, or their interplay, may contribute to knotweed invasion success (Murrell et al., 2011; Parepa et al., 2013; Siemens & Blossey, 2007). However, the precise mechanisms and in particular microbial taxa involved in these phenomena are unknown.

In a preliminary screening of some invasive knotweed populations around Tübingen (see Supplementary Information) we had found a large diversity of root-associated fungi, and 40% of the studied fine-root samples also harboured ITS sequences of Sebacinales fungi, including *S. herbamans* (Table S1). Thus, interactions between invasive knotweed and Sebacinales appeared to be common, and we were curious about the nature of the interaction between the two taxa, in particular whether these hidden and little understood, but very common, fungi influenced the growth and performance of invasive knotweed. We suspected that if *S. herbamans* had an effect on knotweed, it would depend on environmental conditions, in particular the resource supply of the plants. We tested this through a greenhouse experiment in which we inoculated Japanese knotweed and its hybrid with *S. herbamans* under benign as well as drought, shading and low-nutrient conditions. To confirm inoculation success and quantify fungal colonization, we used immunofluorescence microscopy and qPCR. Specifically, we asked the following questions: (1) Does colonization of knotweed by *S. herbamans* depend on environmental conditions? (2) What effects does the endophyte have on the growth and performance of knotweed, and (3) to what extent are these effects environment-dependent?

## Material and Methods

### Plant material

*Reynoutria japonica* and its hybrid *Reynoutria* × *bohemica* are large perennial forbs from the Polygonaceae family that have become invasive in riparian and ruderal habitats in the temperate regions of Europe and North America. They are clonal plants with extensive rhizome networks, and in their invasive range they often form dense monoclonal stands and become extremely dominant (Aguilera et al., 2010). In our experiment, we used plant material from a live collection of knotweed clones that had originally been collected across seven regions in Switzerland and Germany (Krebs et al., 2010) and that had been cultivated in a common garden for several years. We used rhizome cuttings from 20 *R. japonica* clones and 13 *R.* × *bohemica* clones, with approximately ten rhizome fragments, each containing two nodes and thus one intact internode, from each clone. After removal of all fine roots, the rhizome fragments were surface-sterilized using the method described by (Huang et al., 2014).

### Endophyte material

We worked with the endophyte *Serendipita herbamans* (DSM 27534), a member of the order Sebacinales, whose discovery and isolation was described in (Riess et al., 2014). Prior to the experiment, we grew the endophyte for 14 days in Petri dishes with MEA medium containing 2% malt extract and 1.5% agar at 20°C in the dark. We then used 5 mm plugs from these plates, containing media and mycelia, to inoculate 0.5 L Erlenmeyer flasks with 250 ml malt extract (2.0%) liquid medium. The inoculated flasks were incubated in the dark on a rotary shaker (47-52 rpm) at 20°C. After two weeks of incubation, the resulting mycelium was separated from the media and washed five times with sterile distilled water.

### Experimental set-up

We set up a greenhouse experiment in which we tested the effects of endophyte inoculation on knotweed growth in four different environments: control, drought, low nutrients and shade. Except for the control environment, all conditions were expected to be stressful for the plants. We planted individual rhizome fragments 3 cm deep in 1.5 L pots filled with a 1:3 mixture of low-nutrient field soil and sand (Sand- und Kieswerk Bischoff, Rottenburg, Germany). Prior to planting, we measured the length and diameter of each rhizome fragment. All pots were placed on individual saucers and watered as needed. After two weeks, when all aboveground shoots had appeared, we inoculated half of the pots with 0.5 g of fresh *S. herbamans* mycelium which were applied in small pits close to the center of the rhizomes (Fig.1A). For non-inoculated plants we also created the same pits and applied a similar volume of distilled water. Another two weeks after the inoculation, we started the environmental treatments. In the shade treatment, the plants were covered individually with shading mesh bags that reduced light levels to approximately 20%. The low-nutrient plants did not receive any fertilizer throughout the experiment, whereas all others received 7:3:6 N:P:K fertilizer (b1 Universal-Flüssigdünger, toom Baumarkt GmbH) equivalent to 150 kg N/ha distributed across 15 applications at seven-day intervals during the stress treatments. The drought plants generally received only a third of the regular watering amount and, in contrast to all other plants, regularly showed signs of wilting (loss of turgor). A total of 288 plants (160 *R. japonica* and 128 *R.* × *bohemica*) were randomly assigned to the eight treatment combinations (four environmental conditions, with or without endophytes), with approximately equal representation of the two taxa in each treatment. Throughout the experiment, the plants were grown in a climate-controlled greenhouse, in a completely randomized order, with supplemental lighting at a 14:10 h light:dark cycle at 20°C/18°C.

**Figure 1.**
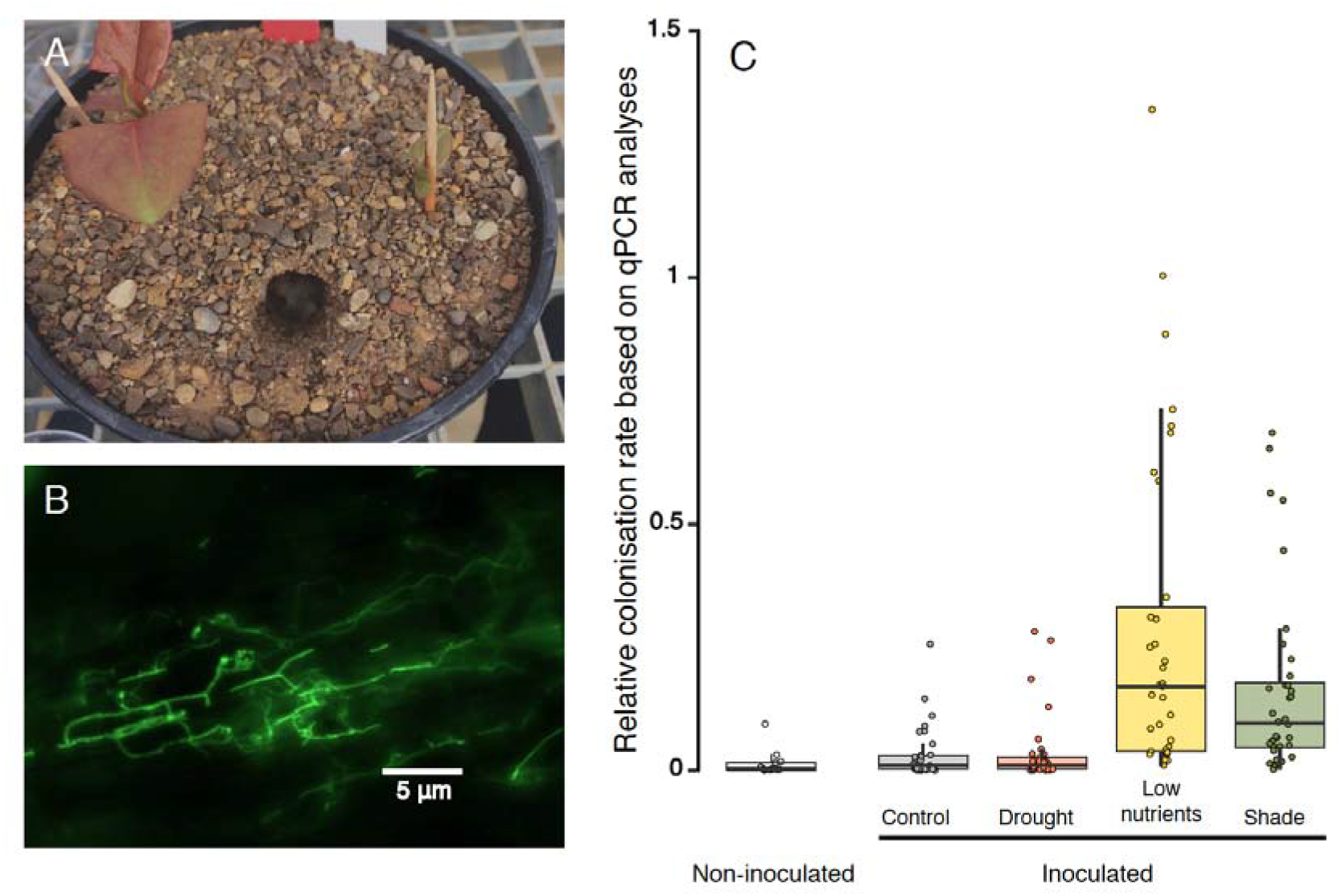
Experimental inoculation of *Reynoutria* plants with *Serendipita herbamans*, and the resulting fungal colonization. (A) An experimental pot right after inoculation, with freshly regenerated knotweed and the pit through which *S. herbamans* mycelium was added. (B) Fluorescence microscopic image of a root section of an inoculated *Reynoutria* plant, stained with WGA-AF 488. (C) Relative colonization rates of *S. herbamans* in the different experimental treatments, based on qPCR analyses. Points = individual observations; boxes = 25th - 75th percentiles; thick horizontal lines= medians; whiskers = 10th - 90th percentiles.

### Data collection

15 weeks after the start of the treatments, we measured leaf chlorophyll content on four leaves per plant using a handheld chlorophyll meter (SPAD 502Plus, Konica Minolta, Osaka, Japan). We then removed all leaves from the stem, measured their area with a LI-3100C leaf area meter (LI-COR Environmental, Lincoln, Nebraska, USA) and dried the leaves and stems of each plant separately at 80◻ for 3 days. We used the leaf area and leaf dry mass to calculate the specific leaf area (SLA) of each plant, and we combined the leaf and stem dry mass to total aboveground biomass. Finally, we carefully washed the roots of each plant and took 10 fine root samples from different parts of the root system that we mixed and immediately stored at −20°C for subsequent DNA extraction, or placed in fixing solution (0.15% (wt/vol) trichloroacetic acid in 4:1 (vol/vol) ethanol/chloroform) for microscopy.

### Assessment of endophyte colonization

We assessed the fungal colonization of knotweed roots qualitatively through fluorescence microscopy and quantitatively through qPCR. Both analyses were done for randomly selected subsets of the plants. For the microscopy, we collected roots from three pots of each treatment by species combination, including both inoculated and non-inoculated samples (altogether 16 × 3 = 48 samples). The root samples were stained with Wheat Germ Agglutinin-Alexa Fluor 488 (WGA488; Thermo Fisher, Waltham MA, USA), which specifically stains fungal cell walls. The staining procedure was as described in (Deshmukh et al., 2006), and the images were recorded on a Leica TCS SP5 2 confocal microscope using the bright field channel and a GFP filter set for detection of WGA488.

For the qPCR, we analysed 10 plants from each treatment by species combination, i.e. a total of 80 plants, with three replicates of non-inoculated plants and seven replicates for inoculated plants. We ground fine roots to a fine powder in liquid nitrogen using a sterile mortar and pestle, and we used 500 mg of this material to isolate DNA using the DNAeasy Plant Mini Kit (Qiagen, Hilden, Germany). The relative amounts of *S. herbamans* DNA in the samples were determined through qPCR reactions with a *S. herbamans*-specific and a *Reynoutria*-specific primer pair. The *S. herbamans-*specific primer pair SerhaITS binds to the ITS region of the 5.8 S rDNA sequence of *S. herbamans* (SerhaITSfw199: 5’-AGCCTTGTGCGGTAAAGCGA-3’, SerhaITSrev199: 5’-TGTATTCCGGCACCTTAACCTC-3’). The *Reynoutria*-specific primer pair FallCHS binds to the genomic DNA of the Chalcone synthase gene EF090266.2 of *Fallopia japonica* (now *Reynoutria japonica*) (FallCHSfwd: 5’-GGAGATGCGTGTATATTCTT-3’, FallCHSrev: 5’-CCAAAGATGAAGCCATGTAG-3’. The PCR primers were designed using Primer-BLAST (Ye et al. 2012). For the PCR amplification we ran real-time PCRs on a Biorad CFX96 Thermocycler (BioRad, Hercules CA, USA) using the ABsolute SYBR Capillary Mix (Thermo Fisher, Waltham MA, USA) in a final volume of 20 μl, and the following cycler programmes: 95°C for 15 min followed by 45 cycles of 95°C for 15 s, 55°C for 20 s and 72° C for 20 s for the FallCHS primer pair, and 95°C for 15 min followed by 45 cycles of 95°C for 15 s, 60°C for 15 s and 72°C for 10 s for the Serha199 primer pair. To calculate relative amounts of *S. herbamans* DNA, we used the 2^−deltaCt^ method (Livak & Schmittgen, 2001) using the raw threshold cycle (Ct) values determined for the *S. herbamans*- and the *Reynoutria*-specific primer pairs.

### Statistical analyses

To test for species differences in, and effects of environmental conditions on endophyte colonization, we analysed the relative *S. herbamans* densities, as determined by qPCR, with a linear model that included the effects of *Reynoutria* species, environmental treatment, and their interaction as fixed factors. We analysed knotweed responses to endophytes and environments with regard to three variables: aboveground biomass, leaf chlorophyll content, and specific leaf area. For each response variable we fitted a linear mixed model with fungal inoculation, environmental treatments, knotweed species, and their interactions included as fixed factors, and clone identity included as random factor. To account for possible influences of initial size differences, we included the volume of the planted rhizome as a covariate in all three analyses. Prior to the analyses, the biomass and specific leaf area data were log-transformed to achieve homoscedasticity. All linear models were fitted with the *lmer* function in the *lme4* package (Bates et al., 2015) in R (R Core Team, 2018). We used the *effects* (Fox, 2003) and *ggplot2* (Wickham, 2009) packages to visualize results.

## Results

The experimental inoculation of knotweed plants with *Serendipita herbamans* was successful, but relative colonization rates were strongly environment-dependent (Fig. 1C; main effect of environmental treatment in the linear model: *F* = 27.12, *P* < 0.001). While there were hardly any fungi present in the non-inoculated samples, and the average relative colonization levels remained low in the inoculated control and drought treatments, colonization increased four-fold and eight-fold, respectively, under shaded and low nutrient conditions (Fig. 1C). There were no differences among the two *Reynoutria* species in terms of relative fungal colonisation (*F* < 1 and *P* > 0.5 for species main effect and species × treatment interaction). The colonization of *Reynoutria* roots by *S. herbamans* was confirmed by fluorescence microscopy. In all root samples from inoculated plants we detected hyphal structures typical for *S. herbamans* on root surfaces, between the outer cell layers, and inside of some cortical root cells (Fig. 1B). We also observed some hyphal structures in the roots of non-inoculated plants.

As expected, the stress treatments in our experiment strongly impacted the growth of knotweed (Table 1, Fig. 2). Compared to control plants, the biomass was reduced in all three stress treatments, but particularly strongly under low-nutrient conditions. There were also strong treatment effects on chlorophyll content and SLA, with a particularly low chlorophyll content at low nutrient availability, and the highest SLA under shaded conditions (Fig. 2). There were also differences between the two knotweed taxa (Table 1). The hybrid *Reynoutria* × *bohemica* was generally larger (+ 25%) and had a higher SLA (+ 5%) than *R. japonica*, and its biomass was less sensitive to drought and shading than that of *R. japonica* (Fig. S1).

**Table 1.** Analysis of variance testing the effects of inoculation with *Serendipita herbamans*, stress treatment and knotweed species (*Reynoutria japonica* or *R.* × *bohemica*), and their interactions, on the performance of invasive knotweed. Each linear mixed model additionally included the volume of the planted rhizome as a covariate, as well as knotweed clone identity as a random variable. Significant *P*-values are in bold.

|  | df | Aboveground biomass |  | Chlorophyll content |  | Specific leaf area |  |
| --- | --- | --- | --- | --- | --- | --- | --- |
|  |  | *F*-ratio | *P*-value | *F*-ratio | *P*-value | *F*-ratio | *P*-value |
| Rhizome volume | 1 | 12.65 | **<0.001** | 4.32 | **0.038** | 0.51 | 0.474 |
| Fungus | 1 | 2.11 | 0.146 | 0.19 | 0.661 | 0.01 | 0.933 |
| Treatment | 3 | 2111.23 | **<0.001** | 382.03 | **<0.001** | 894.68 | **<0.001** |
| Species | 1 | 13.44 | **0.001** | 3.02 | 0.089 | 7.51 | **0.009** |
| Fungus x Treatment | 3 | 2.65 | **0.049** | 6.56 | **<0.001** | 2.20 | 0.088 |
| Fungus x Species | 1 | 0.24 | 0.624 | 0.56 | 0.454 | 0.72 | 0.397 |
| Treatment x Species | 3 | 3.83 | **0.010** | 7.81 | 0.525 | 1.33 | 0.265 |
| Fungus x Treatment x Species | 3 | 0.31 | 0.817 | 1.16 | 0.324 | 0.29 | 0.829 |
| # Observations |  | 283 |  | 283 |  | 281 |  |

**Figure 2.**
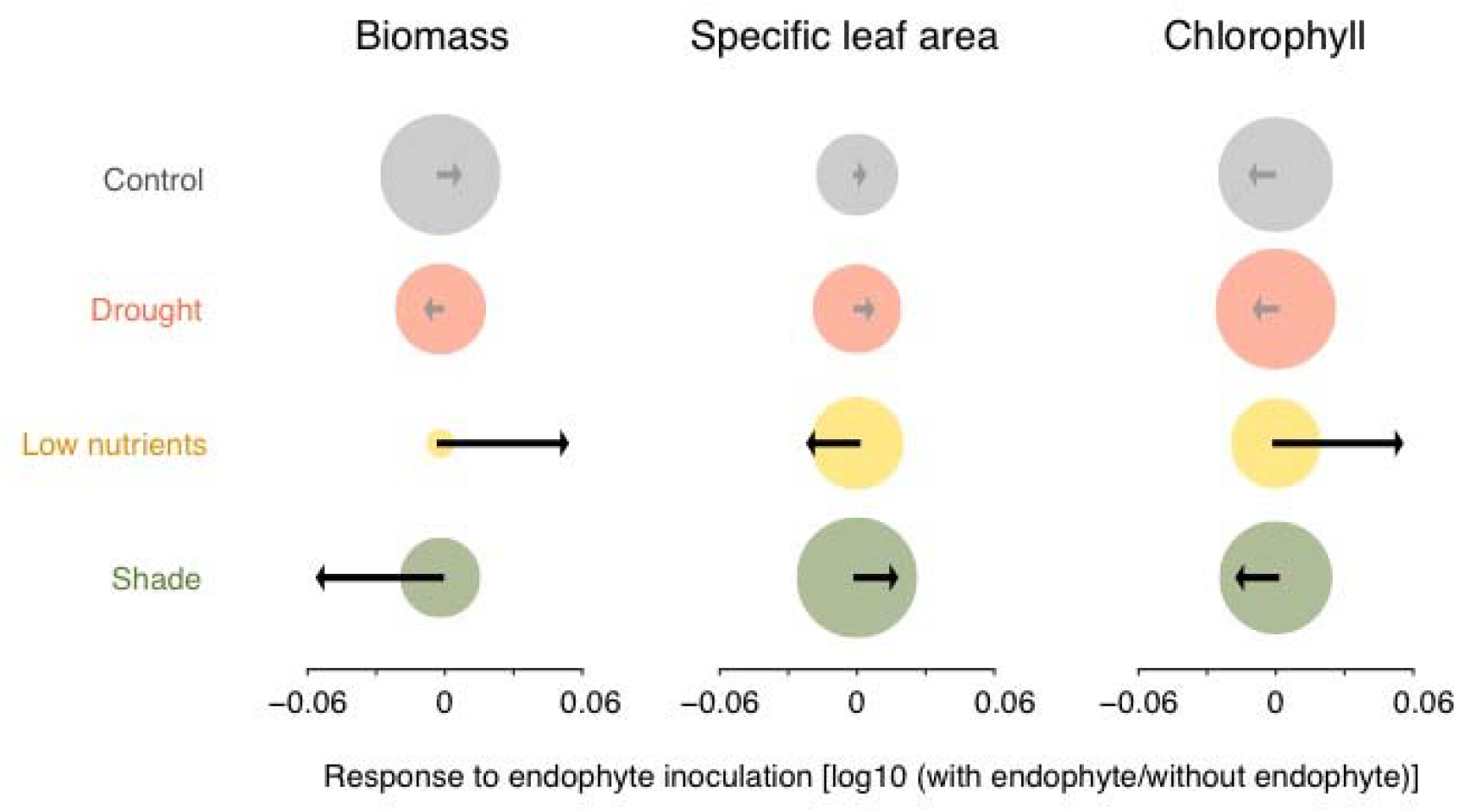
Effects of inoculation with *Serendipita herbamans* on invasive knotweed aboveground biomass, leaf chlorophyll content, and specific leaf area under different types of stress and in control conditions. The disc areas are proportional to the estimated means without endophyte inoculation, and the arrows show the log-responses of plants to endophyte inoculation in each treatment. Black arrows represent significant endophyte effects within a particular treatment, as indicated by Tukey’s HSD test, grey arrows are non-significant.

Inoculation with *Serendipita herbamans* influenced the growth of the knotweed plants, but again in a strongly treatment-dependent manner, with significant fungus by treatment interactions (but no fungus main effects) for aboveground biomass and chlorophyll content, and a marginally significant interaction for SLA (Table 1, Fig. 2). Under low-nutrient conditions, addition of the endophyte increased knotweed biomass by 15%, but decreased it by 10% in the shade, or had no effect at all under drought or control conditions. Similarly, endophyte inoculation increased chlorophyll content by 13% under low nutrients but decreased it by 5% under shaded conditions, and had no significant effects in the other two treatments. Finally, the SLA was positively affected in the shade but negatively under low-nutrient conditions, with no effects under drought or control conditions.

## Discussion

Plant-microbe interactions play an important role in natural ecosystems (Bever et al., 2012; Klironomos, 2002; van der Putten et al., 2013). However, the ecological function of endophytic microbes that live within plants is so far little understood. In this study we show that the fungal root endophyte *Serendipita herbamans* can rapidly colonize invasive knotweed (*Reynoutria* ssp.) and influence its growth, with detrimental effects in the shade but promotion of growth under low-nutrient conditions. Our study thus demonstrates that this widespread endophyte interacts with an important invasive plant, and it also highlights the environment-dependency of plant-endophyte interactions.

### Endophyte colonization

Compared to previous studies on plant-Sebacinales interactions, our experiment had a rather realistic set-up, with fungi inoculated into a non-sterile natural soil that presumably already contained a microbial community. Microscopy and qPCR confirmed that our inoculations were successful and that *S. herbamans* was able to colonize knotweed plants, which confirms field observations in the Tübingen area where Sebacinales including *S. herbamans* are frequent endophytes of invasive knotweed populations (Table S1). This is not surprising, given the broad host range of *Serendipita herbamans*, and of Sebacinales in general, which also includes native Polygonaceae (Garnica et al., 2013; Riess et al., 2014). Although exotic species are known to lose specialised biotic interactions, they often interact with generalist enemies and mutualists in the introduced range (Mitchell & Power, 2003; Richardson et al., 2007; van Kleunen et al., 2018). However, so far we do not know how novel the interaction between knotweed and *S. herbamans* really is because there are no data on endophyte diversity from the native East Asian range.

We found that the relative colonization of knotweed plants by *S. herbamans* was generally much stronger under low-nutrients or shade conditions than under control or drought conditions. Thus, *Reynoutria* plants appear to actively regulate their interactions with *S. herbamans* in an environment-specific fashion. It is known that plants can control fungal colonization, e.g. through the production of defense compounds or secondary metabolites inhibiting microbial growth (Zipfel & Oldroyd, 2017), or by diverting more carbohydrates to fungal symbionts (Carbonnel & Gutjahr, 2014; Martin et al., 2017). This has also been shown for the closely related *S. indica* which interferes with the immune system of host plants (Jacobs et al., 2011) and influences sugar concentrations in their roots (Opitz et al., 2021). The functional and adaptive explanation for this is usually that plant benefits from interactions with fungi are environment-dependent, and therefore plants stimulate or restrict fungal access depending on these benefits. For instance, mycorrhizal colonisation is often triggered by low-nutrient conditions (Bueno de Mesquita et al., 2018). We also found that relative fungal colonization was highest under low-nutrient conditions, which is in line with the idea that *S. herbamans* improves the nutrition of *Reynoutria* plants. It is less clear why relative colonization was also increased in shaded plants because these should have been mainly carbon-limited, and under such conditions plant-microbe interactions often turn parasitic, as has been shown e.g. for interactions with mycorrhiza or rhizobia (Ballhorn et al., 2016; Lau et al., 2012).

We also detected *S. herbamans* in some non-inoculated plants. The sources of this could be external, e.g. fungi spores present in the potting soil, or splash dispersal from adjacent pots. However, the most likely explanation seems that *S. herbamans* was already present in some of the planted rhizomes. We know that some invasive knotweed populations are naturally colonized by *S. herbamans*, and we therefore cannot rule out that some surface-sterilized rhizomes still harboured the fungus.

### Endophyte effects on plant growth

The inoculated *S. herbamans* fungi not only successfully colonized the knotweed plants in our experiment, but they also significantly impacted their growth. The magnitude and direction of these effects were strongly environment-dependent. Under benign or drought conditions, endophyte effects on plants were small and non-significant, whereas under low-nutrient conditions inoculation had strong positive effects, and under shade conditions strong negative effects on knotweed performance. Similar context-dependent effects of endophytes have been found in other study systems (Davitt et al., 2010; Laitinen et al., 2016; Shaffer et al., 2018).

Low-nutrient conditions greatly reduced knotweed biomass, and here relative *S. herbamans* colonization was strong and the fungus increased plant growth. This suggests an active promotion of endophyte access by the plants because the fungi improve plant nutrition under these conditions. The observed increase of leaf chlorophyll content, which strongly correlates with leaf nitrogen content (Evans, 1989), supports this idea. We know that *S. herbamans* improves plant growth under lab conditions (Riess et al., 2014), and that the closely related *Serendipita indica* can improve the nutrient acquisition and growth of many plant species (Achatz et al., 2010; Barazani et al., 2005; Giauque et al., 2019; Varma et al., 1999; Waller et al., 2005). Thus, it seems very likely that *S. herbamans* also improved the nutrition, and as a consequence biomass growth, of invasive knotweed in our experiment.

Under shade conditions, the effects of *S. herbamans* were reversed, and inoculation negatively affected knotweed biomass as well as leaf chlorophyll content, suggesting that under these conditions the fungus indeed turned parasitic and compromised plant nutrition. Similar shifts in the directions of plant-microbe interactions have been observed in other studies (Ballhorn et al., 2016; Lau et al., 2012), and the likely explanation is that the typical ‘trade logic’ of mutualistic plant-microbe interactions-microbes receive photosynthates in exchange for improved nutrient uptake - only works where soil nutrients are limiting, but under carbon-limited shade conditions, it does not.

In the control and drought treatments, colonization and growth effects of endophytes were very low, indicating that under these conditions the host plants limited fungi access, similar to what is known from plant-mycorrhiza interactions (Averill et al., 2019; Carbonnel & Gutjahr, 2014). For the drought treatment, with episodes of plant wilting, it is also possible that the spread of fungi was simply limited by the lack of moisture.

Our results that *S. herbamans* can promote or weaken knotweed growth depending on environmental context raises intriguing questions about the habitat preferences of invasive knotweeds. Across their invasive range in Europe and North America, the species mostly thrive in open and nutrient-rich habitats, and benefit in particular from fluctuating nutrient supply (Parepa, Fischer, et al., 2013), but they rarely spread under closed canopy (Beerling, 1991; Pyšek et al., 2009). It is possible that interactions with *S. herbamans* or other microbes contribute to these habitat preferences, by facilitating nutrient uptake in open habitats but limiting knotweed under shaded conditions. Further research - in particular field experiments - is needed to test these hypotheses.

### Conclusions

Our study demonstrates that the common fungal endophyte *Serendipita herbamans* can rapidly colonize fine roots of invasive knotweed and influence its growth both positively or negatively, depending on the environmental context. As *S. herbamans* is present in at least some invasive knotweed populations, the fungus could play a role in the growth and success of knotweed in some invaded habitats. However, understanding the true significance of this plant-fungus interaction requires further data, because ecological communities are of course more complex than our experimental set-up. In its natural habitat, invasive knotweed also interacts with competitors, herbivores and other enemies and mutualists, and some of these might be interacting with *S. herbamans*, too. Thus, the next step should be multi-species experiments, in the field or using mesocosm approaches, that evaluate the impact of *S. herbamans* on knotweed and other plants in a community context.

## Acknowledgments

We thank Christiane Karasch-Wittmann, Sabine Silberhorn, Julia Rafalski, Mirjam Rieger, Mohamed Osman, and several student helpers for their assistance with the greenhouse experiment and/or molecular work. This project was supported by the German Research Foundation (DFG grant BO 3241/8-1 to OB), and by a scholarship of the China Scholarship Council (grant no. 201504910498) to Zhiyong Liao.

## Authors’ contributions

SG, MP and OB conceived and designed the experiment, SG, ZL and SH carried out the experiment, FW performed microscopy and qPCR analyses, MP and OB analyzed the data, SG, FW, MP and OB drafted the manuscript, and all authors contributed to its revision.

## Data availability

All data from our experiment will be made available through Dryad. The sequencing data from the supplement are stored at GeneBank under MZ650923 - MZ651047.

