## Supplemental Data 1 for "Environmental stress determines the colonization and impact of an endophytic fungus on invasive knotweed" (from another version of this preprint: version 1, 2021-09-11)

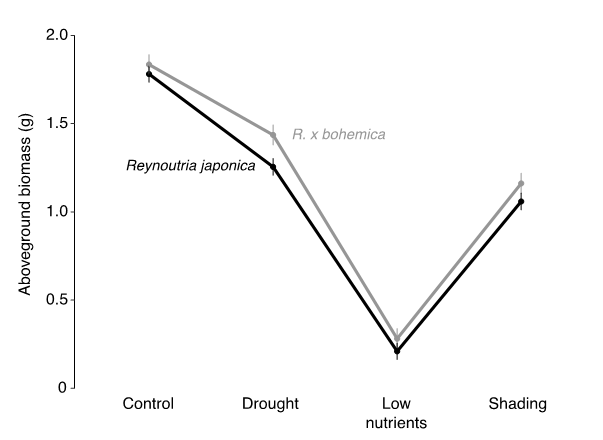

**Fig. S1.** Biomass responses to the stress treatments, separately for the two knotweed species (the significant Treatment x Species interaction in Table 1). The data are means and standard errors estimated from the linear mixed model.

**Methods for Table S1**

*Sampling sites*

We collected roots and rhizomes of *Reynoutria japonica* Houtt. in September 2013 from six sites around Tübingen, Germany, three close to a river (A-C) and three from ruderal habitats (D-F). The riverside sites were: (A) Pliezhausen I (48°33'20.3"N, 9°11'28.6"E), with a knotweed population size of approximately 200 m^2^, 10 m distance to the next ectomycorrhizal tree (*Carpinus betulus*), and a vegetation dominated by *Acer* sp., *Euonymus europaeus*, *Salix* sp., and *Vicia sepium*. (B) Pliezhausen II (48°33'19.1"N, 9°11'26.5"E), with a population size of 4 m^2^, 5 m distance to the next *Carpinus* tree, and vegetation dominated by *Acer campestre*, *Alnus* sp., *Carpinus betulus*, *Cornus sanguinea*, *Equisetum telmateia*, *Euonymus europaeus* and *Salix* sp. (C) Rottenburg I (48°27'55.2"N, 8°55'00.3"E), with a population size of 500 m^2^, 2 m to the next ecomycorrhizal tree (*Salix* sp.), and vegetation dominated by *Alnus* sp., *Corylus avellana*, *Humulus lupulus*, *Picea abies*, *Prunus* sp., *Salix* sp. and *Sambucus nigra*. The ruderal sites were: (D) Rottenburg II (48°28'14.9"N, 8°56'56.7"E), with a population size of only 2 m^2^, 8 m to the next *Carpinus* tree, and vegetation dominated by *Acer campestre*, *Carpinus betulus*, *Cornus sanguinea*, *Euonymus europaeus* and *Quercus* sp. (E) Tübingen I (48°30'30.9"N, 9°02'31.9"E), with a population size of 40 m^2^, 15 m distance to the next *Carpinus* tree, and vegetation dominated by *Acer campestre*, *Acer pseudoplatanus* and *Robinia pseudoacacia*. (F) Tübingen II (48°32'19.5"N, 9°01'47.5"E), with a population size of 3 m^2^, 4 m to the next ectomycorrhizal *Pinus sylvestris* tree, and vegetation dominated by *Acer* sp., *Carpinus betulus*, *Malus domestica*, *Pinus sylvestris*, *Prunus* sp. and *Quercus robur*.

*Molecular lab work*

We ground each dried root and rhizome sample with a sterile plastic pestle in liquid N for several times and then extracted DNA with the innuPREP Plant DNA Kit (Analytic Jena AG, Jena, Germany), following the manufacturer's protocol. To detect root-associated fungi, we ran PCRs using the universal fungal-specific primers ITS1F (Gardes and Bruns, 1993) and ITS4 (White et al., 1990), and MangoTaq^TM^ DNA Polymerase (Bioline GmbH, Luckenwalde, Germany). The PCRs had 25 μl reactions containing 14.5 μl of dd H_2_O, 5 μl of 5 x Reaction Buffer coloured, 0.75 μl MgCl_2_ (50 mM), 1 μl dNTPs (5 mM), 0.5 μl forward primer (25 pmol/μl), 0.5 μl reverse primer (25 pmol/μl), 0.25 μl Mango-Taq DNA Polymerase (2 u/μl) and 2.5 μl DNA template. The PCR amplification conditions were: 3 min initial denaturation at 94°C, then 10 x (30 s denaturation at 94°C, 45 s annealing at 60°C - 1°C per loop, 75 s elongation at 72°C), 26 x (30 s denaturation at 94°C, 45 s annealing at 50°C, 75 s elongation at 72°C), and 7 min at 72°C. To detect Sebacinales, we did a second round of PCRs with the Sebacinales-specific primers NSseb1 (Weiss et al., 2011) and NL4 (White et al., 1990), and Phusion^®^High-Fidelity DNA Polymerase (Thermo Fisher Scientific, Schwerte, Germany). The previous PCR reactions were used for a nested PCR with the Sebacinales specific primer NSseb2 (Garnica et al., 2013) and NL4 and Phusion Polymerase. The PCRs were in 25 μl reaction volumes containing 3 μl dd H_2_O, 5 μl GC-buffer 5 x (incl. 7.5 mM MgCl_2_), 1 ul dNTPs (5 mM), 0.5 μl forward primer (25 pmol/μl), 0.5 μl reverse primer (25 pmol/μl), 0.25 μl Phusion Polymerase (2 u/μl) and 2.5 μl DNA template. The PCR amplification included an initial denaturation at 98°C for 30 s and 34 x (10 s denaturation at 98°C, 20 s annealing at 64°C, 30 s elongation at 72°C), and 10 min at 72°C. Single-band PCR products were sequenced directly as described below.

We verified PCR results with agarose gel electrophoresis (0.7% agarose with 1 x TBE buffer). The PCR products were cloned using the TOPO^®^ ^TA^ Cloning^®^ Kit (Invitrogen^TM^, LifeTechnologies^TM^ GmbH, Darmstadt, Germany). The ligation premix for eight samples consisted of 2 μl Salt solution (1.2 M NaCl, 0.06 M MgCL_2_), 4 μl dd H_2_O and 2 μl PCR^TM^ 4-TOPO^®^TA vector. For each sample 1 μl DNA and 1 μl ligation premix was mixed and incubated for 30 min at room temperature. Then 15 μl of One Shot^®^ Chemically Competent TOP 10 *Escherichia coli* cells were added, and after short centrifugation the samples were incubated for 30 min on ice. Thereafter we placed the samples in a water bath at 42°C for 30 s, transferred them to ice for 2 min, added 80 μl of S.O.C. medium to each sample and shake-incubated it for 1 h at 200 rpm and 37°C. We grew the clones on LB-medium with Kanamycin (1 ml/l) at 37°C over night and subjected at least 12 bacterial colonies from each sample to a PCR with M13 forward and reverse primers (Invitrogen^TM^, Life Technologies^TM^ GmbH, Darmstadt, Germany) and MangoTaq Polymerase. For amplification we used a program with 7 min at 94°C, 32 x (20 s denaturation at 94°C, 40 s annealing at 55°C, 160 s elongation at 72°C), and 7 min at 72°C.

We digested eight clones per sample with HinfI (New England Biolabs GmbH, Frankfurt am Main, Germany). The gel electrophoresis after restriction was conducted with 1% agarose and 10 x SBS buffer. All restriction fragment length polymorphisms of the internal transcribed spacer region (ITS-RLFP) with different restriction patterns and some ITS-RFLPs showing similar phenotypes were sequenced. PCR products were purified with ExoSAP-IT^®^ (Affymetrix UK Ltd, Bucks, UK) diluted 1 : 20 according to the manufacturer's instruction. ABI cycling was conducted with BigDye^®^ Terminator v3.1 Cycle Sequencing Kit (Applied Biosystems^®^, Life Technologies TM GmbH, Darmstadt) using 4 μl ABI PRISM BigDye (1 : 6) and 1 μl M13R primer (12.5 pmol/μl) and if necessary M13F primer (12.5 pmol/μl); Sebacinales: ITS1 (White et al., 1990), ITS4 (White et al., 1990), NLMW1 (Sampaio et al., 2002), NL4 and sometimes ITS2 (White et al., 1990), 5.8SR (Vilgalys and Hester, 1990), LR0R (Vilgalys and Hester, 1990). The cycle sequencing was performed with the program 10 ABI31CYCL: Initial denaturation: 96°C for 1 min, 27 x (denaturation: 96°C for 10 s, annealing: 50°C for 5 s, elongation: 60°C for 4 min). After cycle sequencing, the DNA was precipitated with 75% isopropanol, washed with 80% ethanol, dried, dissolved in HiDi and placed in the ABI3130xl (Applied Biosystems, Foster City, CA, USA) for sequencing.

*Taxonomic assignment of fungi*

We assembled forward and reverse DNA sequences using Sequencher v.4.9 (Gene Codes Corporation, Ann Arbor, MI, USA) and saved them as consensus sequences. To identify root-associated fungi of *R. japonica*, these consensus sequences were submitted to BLAST (Altschul et al., 1990) searches against the rRNA/ITS reference sequences for fungi deposited in the NCBI GenBank ([www.ncbi.nlm.nih.gov](http://www.ncbi.nlm.nih.gov)), with the the *maximum* number of *target sequences* limited to 10.

*References*

Altschul SF, Gish W, Miller W, Myers EW, Lipman DJ (1990) Basic local alignment search tool. J Mol Biol 215: 403-410.

Gardes M, Bruns TD (1993) ITS primers with enhanced specificity for basidiomycetes - application to identification of mycorrhizae and rusts. Mol Ecol 2: 113-118.

Garnica S, Riess K, Bauer R, Oberwinkler F, Weiss M (2013) Phylogenetic diversity and structure of sebacinoid fungi associated with plant communities along an altitudinal gradient. FEMS Microb Ecol 83: 265-278.

Sampaio JP, Weiss M, Gadanho M, Bauer R (2002) New taxa in the Tremellales: *Bulleribasidium oberjochense* gen. et sp. nov., *Papiliotrema bandonii* gen. et sp. nov. and *Fibulobasidium murrhardtense* sp. nov. Mycologia 94: 873-887.

Vilgalys R, Hester M (1990) Rapid genetic identification and mapping of enzymatically amplified ribosomal DNA from several *Cryptococcus* species. *J Bacteriol* 172: 4238-4246.

Weiss M, Sykorová Z, Garnica S, Riess K, Martos F, Krause C, Oberwinkler F, Bauer R Redecker D (2011) Sebacinales everywhere: previously overlooked ubiquitous fungal endophytes. PLoS ONE 6: e16793.

White T, Bruns TD, Lee S, Taylor J (1990) Amplification and direct sequencing of fungal ribosomal RNA genes for phylogenetics. In: Innis et al. (eds) *PCR Protocols: A Guide to Methods and Applications*. Academic Press, pp. 315-322.
